# Better data for better predictions: data curation improves deep learning for sgRNA/Cas9 prediction

**DOI:** 10.1101/2025.06.24.661356

**Authors:** Tyler S. Browne, David R. Edgell, Gregory B. Gloor

## Abstract

The Cas9 enzyme along with a single guide RNA molecule is a modular tool for genetic engineering and has shown effectiveness as a species-specific anti-microbial. The ability to accurately predict on-target cleavage is critical as activity varies by target. Using the sgRNA nucleotide sequence and an activity score, predictive models have been developed with the best performance resulting from deep learning architectures. Prior work has emphasized robust and novel architectures to improve predictive performance. Here, we explore the impact of a data-centric approach through optimization of the input target site adjacent nucleotide sequence length and the use of data filtering for read counts in the control conditions to improve input data utility. Using the existing crisprHAL architecture, we develop crisprHAL Tev, a best-in-class bacterial SpCas9 prediction model with performance that generalizes between related species and across data types. During this process, we also rebuild two prior *E. coli* Cas9 datasets, demonstrating the importance of data quality, and resulting in the production of an improved bacterial eSpCas9 prediction model. The crisprHAL models are available through GitHub (https://github.com/tbrowne/crisprHAL).

## 1 INTRODUCTION

The CRISPR-Cas9 system is capable of a wide range of genetic engineering tasks through the production of DNA double-strand breaks. In this system, the Cas9 nuclease and a single guide RNA (sgRNA) bind a DNA target containing a 20-nucleotide sequence complementary to the target site. The activated nuclease then induces a double-strand DNA break (Deltcheva et al., 2011; Jinek et al., 2012). Binding and cleavage activity by Cas9 requires a protospacer adjacent motif (PAM) immediately downstream of the target DNA. For the commonly used *Streptococcus pyogenes* Cas9 nuclease (SpCas9), this PAM sequence is NGG (Jinek et al., 2012). Along a sequence of DNA this PAM site will occur once every 16 nucleotides on each strand, assuming a random and equal nucleotide (nt) distribution, highlighting the numerous potential targets that exist. Given the ease of designing the sgRNA sequence there is broad potential for the system as a reprogrammable site-specific nuclease. Despite the large number of sites, prior work has shown wide variations in so-called “on-target” activity, defined as cleavage of the chosen target site (Doench et al., 2014; Wang et al., 2014; Doench et al., 2016; Guo et al., 2018; Hamilton et al., 2019; Ham et al., 2023).

Several machine-learning models have been constructed that use the nucleotide sequence of the target site, and often other biological parameters, to predict on-target activity (Guo et al., 2018; Chuai et al., 2018; Wang and Zhang, 2019; Dimauro et al., 2019; Wang et al., 2019; Zhang et al., 2020; Kim et al., 2019; Lin et al., 2020; Wan and Jiang, 2023; Ham et al., 2023). The best performing of these models typically rely upon deep learning architectures to make their predictions. Most existing machine-learning models have been constructed for mammalian applications; only three bacteria-specific Cas9 prediction models exist (Guo et al., 2018; Wang and Zhang, 2019; Ham et al., 2023). Regardless of their application, recent work has generally focused on improving on-target activity predictions through two methods: first by updating the model architecture and second by using datasets with greater numbers of unique sgRNA sequences. While both factors are clearly important, examining and improving the quality of data has been neglected, and this is the aim of this work. We focus on two key aspects to improve predictive performance: improving the accuracy of the on-target activity score, and the inclusion of relevant sequences adjacent to the chosen target site.

On-target sgRNA/Cas9 activity datasets suitable for machine-learning training have been generated using a variety of methods which contrast control and experimental conditions and use DNA sequencing read counts as a proxy for on-target sgRNA/Cas9 activity (Guo et al., 2018; Ham et al., 2023). Due to the compositional nature of these datasets — a single read count is an arbitrary metric dependent upon sequence depth and requires a transformation for interpretability — the read counts are normalized in some fashion before activity scores are calculated (Quinn et al., 2019). The “depletion” datasets that measured killing efficiency of a wild-type and enhanced specificity SpCas9 in *Escherichia coli*, generated by Guo et al. (2018), used a mean normalization followed by a non-targeting control adjustment and then a data standardization using a Z-score statistic. In contrast, the datasets that measured TevSpCas9 and SpCas9 activity in *E. coli* and *Citrobacter rodentium*, generated by Ham et al. (2023), used ALDEx2 (Fernandes et al., 2013) to generate a difference-between metric derived from a centre log ratio-based approach (Fernandes et al., 2014). All datasets measured killing efficiency through genomic cleavage (sgRNA depletion) or elimination of a plasmid encoding a toxin gene resulting in growth (sgRNA enrichment).

Regardless of the method used, low control condition read counts cause two issues. First, low counts reduce the precision of measurement, and second, the absolute range of observable values is smaller (Fernandes et al., 2013; Love et al., 2014; Gloor et al., 2016). In depletion experiments low read counts in the control condition constrain the precisions by which we can measure activity, and in enrichment, inactivity. In either case, this imprecision results in inaccurate activity scores if appropriate filtering for a minimum control condition read count is not performed. Prior to this point, no group has investigated the role that optimizing for the minimal read count in the training set has on model performance. This is not to suggest that dynamic range concerns have been ignored, read count minima cutoffs have been used previously, but these have not been investigaged systematically. We suggest that each dataset may have a minimum read cutoff which balances the reduction in data with the improvement in scoring accuracy.

A second important component to the on-target activity scores is the target site sequence supplied to a model as the input feature. In many existing models, this sequence is simply the 20 nucleotide sgRNA target site, often with the addition of its respective PAM sequence — NGG when using an SpCas9 dataset (Zhang et al., 2020; Chuai et al., 2018; Wang et al., 2019; Dimauro et al., 2019; Wan and Jiang, 2023). However, the nucleotides flanking the target site do contribute to nuclease activity (Wang and Zhang, 2019; Ham et al., 2023). Prior work found Cas9-DNA interactions 8 nucleotides downstream of the sgRNA target site which appear to mediate target search (Yang et al., 2021). More distant interactions were also found 14 nucleotides downstream of the PAM, which appears to regulate nuclease binding and dissociation (Zhang et al., 2019). Work by Ham et al. (2023) examined and optimized model performance through the incremental inclusion of target site adjacent nucleotides to their model. This method found optimal performance using an input sequence comprising the sgRNA, NGG PAM, and 5 nucleotides downstream, noting that the inclusion of upstream nucleotides only decreased performance. Though this provides insight into adjacent nucleotide impacts on activity, the small dataset sizes used potentially constrain the parameter increase required by longer input sequences. The potential for large datasets to compensate for this parameter increase is promising for the capture of more distant target site adjacent nucleotide impacts on activity, provided the data is sufficiently accurate. It should also be noted that different Cas enzymes are expected to have differing nucleotide sequence preferences, especially nucleases with changes in their binding characteristics that increase specificity such as eSpCas9 (Slaymaker et al., 2016). Thus, the input sequence length that is optimal for SpCas9 model performance may be sub-optimal when applied to training an eSpCas9 model, or to an enzyme derived from a different species.

Here we test bacterial Cas9 prediction models constructed on the existing crisprHAL architecture from Ham et al. (2023) to demonstrate the importance of data quality for performance optimization. We present a best-in-class SpCas9 activity prediction model, trained on a *C. rodentium* TevSpCas9 dataset, with performance that generalizes across species. We refer to this new model as crisprHAL_Tev_. We also recreate two prior datasets from Guo et al. (2018) for SpCas9 and eSpCas9 in *E. coli* which demonstrate the general benefit of data optimization on model performance, resulting in the production of an improved bacterial eSpCas9 activity prediction model.

## 2 METHODS

### 2.1 Datasets

For model development, training, and testing we used three large and three small datasets that measured bacterial Cas9 activity by multiple different assays. Table 1 provides information on the experimental design used to generate these datasets and Table 2 provides information on sequencing depth and the number of available single guide RNAs (sgRNA) for training and validation. The three large datasets measure killing efficiency (depletion) targeting sgRNAs to the host genome which activates cleavage when combined with Cas9 expression; in these assays active sgRNAs are depleted from the library. In the small datasets activity is measured through cellular growth (enrichment) when and active sgRNA/Cas9 complex cleaves a plasmid harbouring the CcdB toxin. Our primary SpCas9 prediction model is constructed from a set of 30138 sgRNAs and the read counts from 9 experimental and 1 *E. coli* Epi300 control replicates generated by Ham et al. (2023). This depletion assay measured the activity of sgRNAs in the presence of the dual nuclease TevSpCas9 to cleave the genome of *C. rodentium* and concomitant loss of the active sgRNA from the pool. The three independent testing sets, also created by Ham et al. (2023), were developed in *E. coli* K-12, each with 10 control and 10 experimental replicates. Two of these datasets measured cleavage of a CcdB toxin-harbouring plasmid with the TevSpCas9 and SpCas9 nucleases, referred to as the pTox TevSpCas9 and pTox SpCas9 datasets. The two pTox datasets have overlapping sgRNA target sites, providing the opportunity to compare cleavage by the TevSpCas9 and SpCas9 enzymes directly. An additional testing dataset measured TevSpCas9 cleavage of the KatG gene cloned from *Salmonella enterica* inserted on a plasmid in *E. coli*, and is referred to as the KatG TevSpCas9 dataset.

**Table 1.** Assay design for the six datasets used for training, testing, and benchmarking.

| Dataset Name | Nuclease | Target | Host | Type | Control Condition | Control Replicates | Experimental Replicates | Source |
| --- | --- | --- | --- | --- | --- | --- | --- | --- |
| *C. rodentium* TevSpCas9 | TevSpCas9 | Host genome contig assembly | *C. rodentium* DBS100 | Depletion | *E. coli* Epi300 pre-transformation | 1 | 9 | Ham et al. (2023) |
| *E. coli* eSpCas9 | eSpCas9 | Host genome coding regions | *E. coli* K-12 | Depletion | Catalytically dead eSpCas9 | 2 | 2 | Guo et al. (2018) |
| *E. coli* SpCas9 | Wild-type SpCas9 | Host genome coding regions | *E. coli* K-12 | Depletion | Catalytically dead SpCas9 | 2 | 2 | Guo et al. (2018) |
| pTox TevSpCas9 | TevSpCas9 | Plasmid target (pTox) | *E. coli* K-12 | Enrichment | Repressed expression | 10 | 10 | Ham et al. (2023) |
| pTox SpCas9 | Wild-type SpCas9 | Plasmid target (pTox) | *E. coli* K-12 | Enrichment | Repressed expression | 10 | 10 | Ham et al. (2023) |
| KatG TevSpCas9 | TevSpCas9 | Plasmid host *S. enterica* KatG gene | *E. coli* K-12 | Enrichment | Repressed expression | 10 | 10 | Ham et al. (2023) |

**Table 2.** Dataset composition for the six sgRNA/Cas9 activity datasets used in this paper. *Testing datasets use a fixed control condition minimum read count filtering cutoff of 20. Datasets not used for model training are indicated with a ‘-’ in the training set size column.

| Dataset Name | Initial Pool | Mean Control Read Coverage | Mean Selective Read Coverage | Unfiltered sgRNAs | Filtering Cutoff | Filtered sgRNAs | Training set size | Testing set size |
| --- | --- | --- | --- | --- | --- | --- | --- | --- |
| *C. rodentium* TevSpCas9 | 31596 | 228.6 | 73.2 | 30138 | 56 | 25244 | 20195 | 5049 |
| *E. coli* eSpCas9 | 65928 | 178.5 | 98.7 | 64021 | 31 | 59489 | 47591 | 11898 |
| *E. coli* SpCas9 | 65928 | 106.7 | 49.7 | 61002 | 73 | 33034 | 26427 | 6607 |
| pTox TevSpCas9 | 304 | 1903.9 | 1968.0 | 279 | 20* | 260 | - | 260 |
| pTox SpCas9 | 304 | 261.8 | 321.3 | 303 | 20* | 255 | - | 255 |
| KatG TevSpCas9 | 317 | 1635.8 | 1659.0 | 228 | 20* | 268 | - | 268 |

We also obtained two additional depletion datasets from Guo et al. (2018) that tested a pool of 65928 sgRNAs in *E. coli*. We reconstructed both datasets from publicly available sequencing data and use the original datasets for comparison. For these datasets, *E. coli* eSpCas9 (n=45071) and *E. coli* SpCas9 (n=44163), we use the scores as calculated by Guo et al. (2018) with the 43nt model input sequences from Wang and Zhang (2019), and an improved scoring scheme outlined below. The original datasets and scoring schemes are referred to as the original *E. coli* eSpCas9 dataset and the original *E. coli* SpCas9 dataset.

### 2.2 Expanded eSpCas9 and SpCas9 datasets

For the generation of a new *E. coli* eSpCas9 and a new *E. coli* SpCas9 model, we reconstruct the two datasets from Guo et al. (2018) using sequencing data generated in *E. coli* K-12 substr. MG1655. This reconstruction was performed in order to make use of all available data for model training. The original data was curated based upon a minimum of 5 sgRNAs mapping to an *E. coli* gene for inclusion — valid for their use case, but unnecessary for model training. The initial sgRNA pool comprised 65928 sgRNAs targeting the eSpCas9 or SpCas9 nuclease to the *E. coli* genome. Each dataset consisted of two control and two experimental condition samples, with catalytically dead eSpCas9 or SpCas9 being used for the control condition. The paired-end reads were merged with usearch (Edgar, 2010) and we obtained read counts for sgRNAs from the sequencing output using an in house script, with sequence reads being counted for both replicates in the control and experimental conditions.

### 2.3 Input encoding and activity score generation

The raw read counts for each sgRNA target in control and experimental conditions were used as the input to ALDEx2, with the ALDEx2 diff.btw, a non-parametric log_2_ fold-change metric, used as the activity score; hereafter referred to as log_2_FC (Fernandes et al., 2014). This metric differs from the Z-score activity metric used by Guo et al. (2018) for the original *E. coli* eSpCas9 and SpCas9 datasets used for comparison testing in that it is the expected value of a Bayesian posteriour derived from the data and ensures that all values are linearly related (Aitchison, 1986; Fernandes et al., 2013). The enrichment pTox TevSpCas9, pTox SpCas9, and KatG TevSpCas9 datasets used a control-condition read cutoff of 20. To test control condition read-count minimum cutoff impacts on activity, we constructed the *C. rodentium* TevSpCas9, *E. coli* eSpCas9, and *E. coli* SpCas9 datasets with incrementally increasing minimum average control condition read count cutoffs — 100 variants of each dataset ranging from a cutoff of 1 to a cutoff of 100 (Fig. 1A).

**Figure 1.**
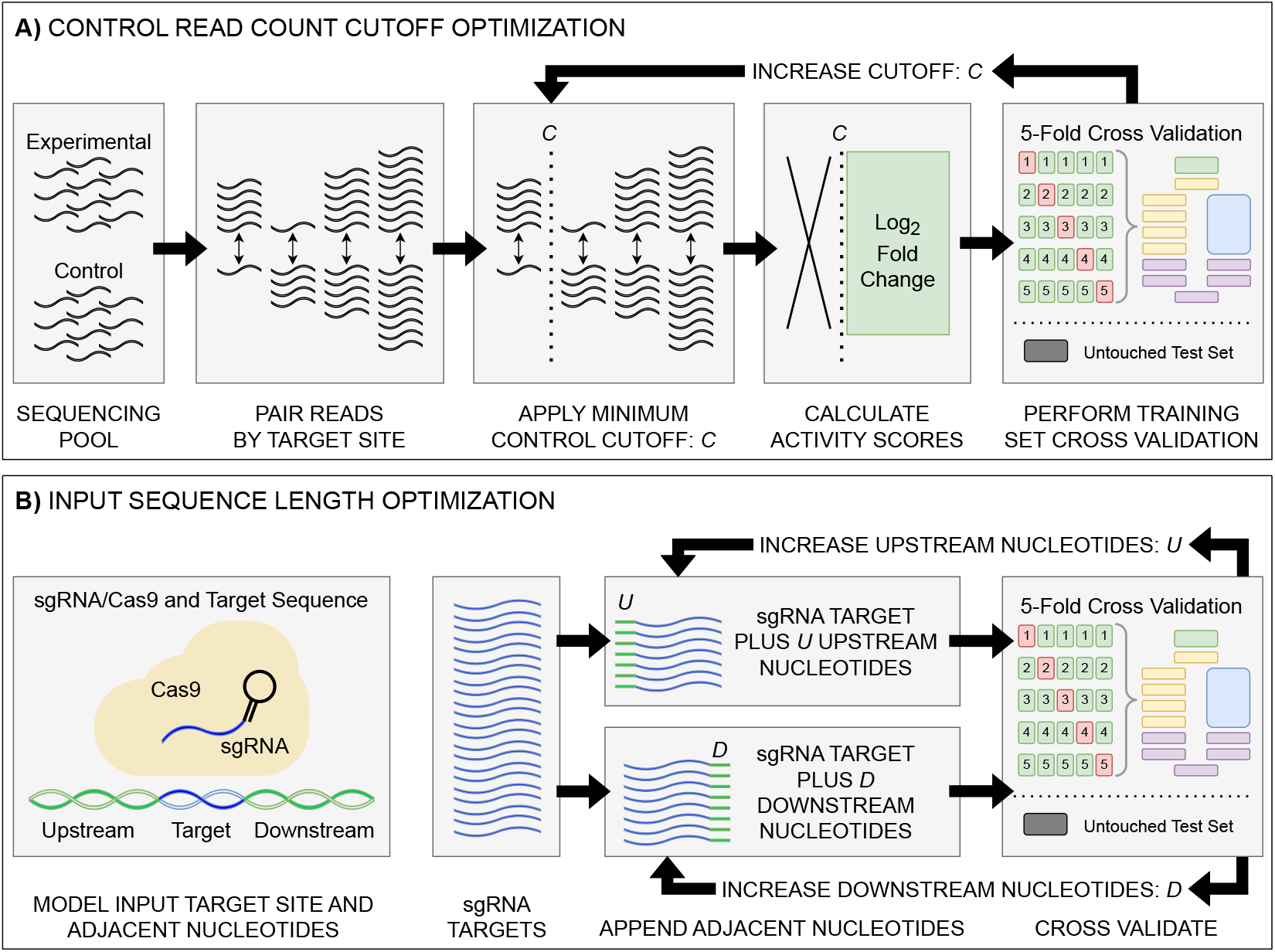
Model input and score optimization workflow. **A)** Identification of an optimal control read count minimum cutoff value *C*. Read counts for each feature in control and experimental replicates were paired by their respective sgRNA target site. Beginning with *C*=1, all sgRNAs with a control read count below the cutoff *C* were removed, with subsequent log_2_FC scores then calculated. The output sgRNAs and corresponding scores were split into training and held-out testing sets. Model performance metrics for the cutoff value were calculated using 5-fold cross validation on the training set. The cutoff value *C* was then increased, with the cutoff curation, score calculation, and 5-fold cross validation repeated. **B)** Identification of optimal upstream and downstream target site adjacent nucleotide inclusion to the model input sequence. Two tests were performed: one upstream and one downstream, starting with values of *U*=0 and *D*=0. Beginning with the 20 nucleotide sgRNA target site, *U* upstream nucleotides were appended, and the data was split into training and testing sets, with subsequent performance metrics obtained via 5-fold cross validation. The same test was performed with *D* downstream nucleotides appended in place of the upstream nucleotides. The process was then repeated, incrementing the number of appended upstream *U* and downstream *D* nucleotides. The optimal input sequence length was then obtained using the optimal *U* and *D* upstream and downstream nucleotides, with a final input length of *U*+20+*D*. Across all optimization tests, like standard model hyper-parameter tuning, the hold-out test set was untouched, being used only to test the final model.

To determine the effect of adjacent nucleotides on prediction accuracy, we extracted the target site sequences from each datasets associated target DNA sequence — *C. rodentium* DBS100, *E. coli* K-12, and the respective plasmid sequences, examining and the impacts to model performance when included alongside the sgRNA and PAM (Fig. 1B). For each set of target sites, we extended the sequence by 500 nucleotide upstream and 150 nucleotide downstream of the sgRNA plus the NGG PAM. Prior to model input, these sequences are transformed into a model-friendly one-hot encoding format of 4×N shape: 4 nucleotide options by an N-length sequence input.

### 2.4 Model architecture

Throughout we use the term “architecture” when referring to the programming setup, and “model” when referring to the output of the architecture after training. Given the focus of this work on improving predictive performance through a data-centric approach, we train all models on the existing crisprHAL model architecture from Ham et al. (2023) (Fig. 2). This architecture uses a deep learning approach with a dual branching structure comprising convolutional and recurrent neural network layers. The convolutional neural network (CNN) layers aim to learn local features of the target site which impact on-target activity through optimizing the weights of convolutional filters, whereas the recurrent neural network (RNN) aims to learn sequential features Rumelhart et al. (1985); LeCun et al. (2015). The dual branching structure passes an encoded sgRNA input through an initial convolutional block to either several successive convolutional blocks or a bidirectional gated recurrent unit (BGRU) RNN (Cho et al., 2014). Each convolutional block consists of a 1-dimensional CNN followed by a LeakyReLU activation function and 1-dimensional max pooling. Each branch then sends their output through two dense layers, each followed by a dropout layer, finally being concatenated into a single output node. This structure is unchanged from the original crisprHAL architecture, save for a small alteration of the BGRU to allow for better GPU usage. A major focus of the original architecture was transfer learning with all parameters frozen in the CNN-branch. Here we do not make use of transfer learning, instead focusing on performance improvement through optimal data curation.

**Figure 2.**
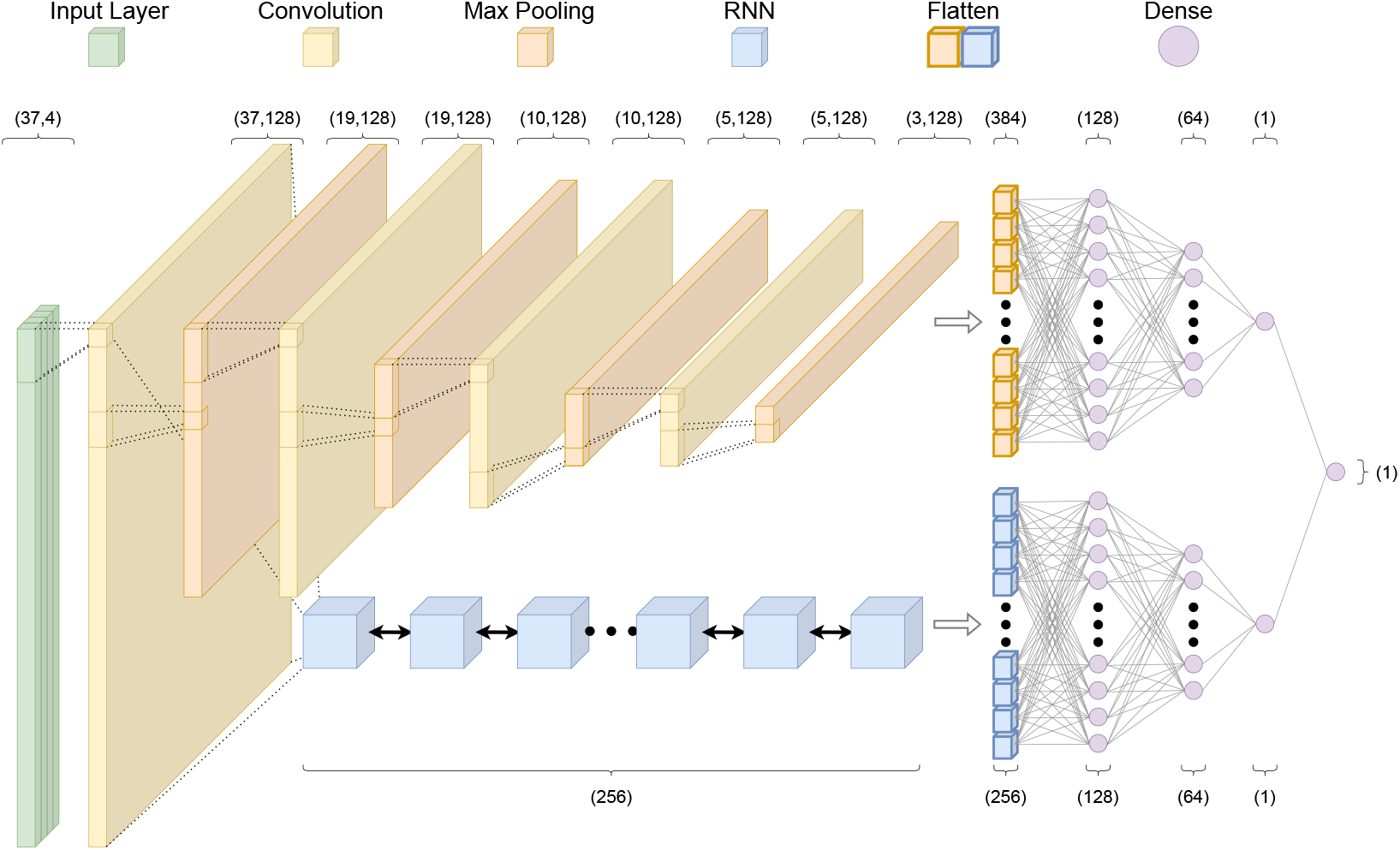
The crisprHAL model architecture. The forward propagation of a one-hot encoded input nucleotide sequence through the dual branch CNN and bi-directional GRU RNN structure, resulting in a final activity prediction. This diagram depicts an input sequence length of 37 nucleotides used by crisprHAL_Tev_.

Hyper-parameter adjustments were made in order to better utilize the larger training sets — the original model was designed for use with small dataset fine-tuning. As such, we reduced the learning rate to 0.0005 and increased the batch size to 1024. Unchanged hyper-parameters include using the “Adam” optimizer with a mean squared error loss function (Kingma and Ba, 2017). For each final presented model, performance is reported following training on the entire training set using hyper-parameters determined by 5-fold cross validation (5CV) using the respective training set. These include the previously mentioned hyper-parameters in addition to the average control condition read count minima (cutoff), upstream and downstream target site adjacent nucleotide inclusion to the model input, and total training epochs.

### 2.5 Prior models

To benchmark our TevSpCas9, eSpCas9, and wild-type SpCas9 models — crisprHAL_Tev_, crisprHAL_eSp_, and crisprHAL_WT_ — we use six existing models from three sources (Table 3). These models are: the eSpCas9 and wild-type SpCas9 models from Guo et al. (2018), Guo_eSp_ and Guo_WT_; the eSp-Cas9 and wild-type SpCas9 models from Wang and Zhang (2019) named here as DeepSgRNA_eSp_ and DeepSgRNA_WT_; and the crisprHAL transfer learning-based TevSpCas9 and wild-type SpCas9 models from Ham et al. (2023), crisprHAL_TL-Tev_ and crisprHAL_TL-WT_. All models are publicly available: crisprHAL (https://github.com/tbrowne5/A-generalizable-Cas9-sgRNA-prediction-model-using-machine-transfer-learning), DeepSgRNA (https://github.com/biomedBit/DeepSgrnaBacteria), and Guo (https://github.com/zhangchonglab/sgRNA-cleavage-activity-prediction). Models have been installed and tested with appropriate configurations as laid out in their respective installation guides. Given the lack of a held out test set for the DeepSgRNA_eSp_ model, crisprHAL_eSp_ model performance reported in this paper is contrasted with their average 5-fold cross validation performance metrics.

**Table 3.** Models compared in this paper. Model name subscript notation is as follows: Tev for TevSpCas9, eSp for eSpCas9, WT for wild-type SpCas9, and the prefix TL for models utilizing transfer learning. *Training data was curated by Guo et al. (2018). †Training data was curated by Ham et al. (2023) with the larger *E. coli* eSpCas9 dataset used for generation of an initial model from which transfer learning was performed with the pTox TevSpCas9 and SpCas9 datasets. We use ‘-’ to denote untested or unreported data and Spearman metrics.

| Model Name | Training Data | Model Design | Activity Scoring | Training CV Set | 5-fold CV Spearman | Held-Out Test Set | Test Set Spearman |
| --- | --- | --- | --- | --- | --- | --- | --- |
| crisprHAL<sub>Tev</sub> | *C. rodentium* TevSpCas9 | CNN-RNN Hybrid | Log<sub>2</sub>FC | 20195 | 0.754 | 5049 | 0.746 |
| crisprHAL<sub>eSp</sub> | *E. coli* eSpCas9 | CNN-RNN Hybrid | Log<sub>2</sub>FC | 47591 | 0.783 | 11898 | 0.783 |
| crisprHAL<sub>WT</sub> | *E. coli* SpCas9 | CNN-RNN Hybrid | Log<sub>2</sub>FC | 26427 | 0.679 | 6607 | 0.684 |
| Guo<sub>eSp</sub> | *E. coli* eSpCas9* | GBR & Bio-feature | Z-Score | 36056 | 0.682 | 9014 | 0.68 |
| Guo<sub>WT</sub> | *E. coli* SpCas9* | GBR & Bio-feature | Z-Score | 35330 | 0.542 | 8833 | 0.54 |
| DeepSgRNA<sub>eSp</sub> | *E. coli* eSpCas9* | CNN & Bio-feature | Z-Score | 45070 | 0.711 | - | - |
| DeepSgRNA<sub>WT</sub> | *E. coli* SpCas9* | CNN & Bio-feature | Z-Score | 44163 | 0.582 | - | - |
| crisprHAL<sub>TL-Tev</sub> | *E. coli* eSpCas9† | CNN-RNN Hybrid | Log<sub>2</sub>FC | 45010 | - | - | - |
|  | pTox TevSpCas9† | & Transfer Learning | Log<sub>2</sub>FC | 279 | 0.630 | - | - |
| crisprHAL<sub>TL-WT</sub> | *E. coli* eSpCas9† | CNN-RNN Hybrid | Log<sub>2</sub>FC | 45010 | - | - | - |
|  | pTox SpCas9† | & Transfer Learning | Log<sub>2</sub>FC | 303 | 0.627 | - | - |

### 2.6 Model performance metrics

To judge model performance we use the Spearman rank correlation coefficient (*ρ*) and Pearson correlation coefficient (*r*); referred to as Spearman correlation and Pearson correlation for simplicity. We report both these metrics to highlight the properties of the predictions since these metrics are concordant in Normally distributed data that is linearly dependent. Across most sgRNA/Cas9 activity prediction models Spearman correlation has been the primary, or only, metric reported, and therefore, this is the main metric we use to benchmark the models. We use the ‘spearmanr’ and ‘pearsonr’ functions from the package SciPy (Virtanen et al., 2020).

## 3 RESULTS

We developed new models for predicting sgRNA activity with SpCas9 trained on depletion datasets by incorporating a different scoring metric, by optimizing the minimum read-count used in each training set, and by including substantially more information from the sequence flanking the sgRNA binding site. We present first the performance of the optimized model in the first section of the results, then show how the individual optimizations improve model performance in the later results sections. In the final results section, we show how these models perform to predict activity in an enrichment experiment which is an independent type of sgRNA assay.

### 3.1 An improved bacterial SpCas9 activity prediction model

For the development of an updated and improved SpCas9 prediction model, we use the hybrid convolutional and recurrent neural network-based crisprHAL architecture from Ham et al. (2023) as a framework. For clarity, we denote the new model generated with the *C. rodentium* dual nuclease-derived TevSpCas9 dataset as crisprHAL_Tev_. The *C. rodentium* TevSpCas9 dataset comprises 25244 sgRNA target-site sequences and their corresponding on-target activity scores from a depletion dataset (Methods) that uses the non-parametric L2FC scoring metric derived from the ALDEx2 tool. Here we used a minimum control read count of 56 (Table 2) derived as in the next section, and included 378 nt input sequence for training, derived as described in the third results section. This dataset was split into 80% training (n=20195) subset, with the remaining 20% subset being used for hold-out testing (n=5049). The training set was used training and model performance optimization (Table 3). Hyper-parameters for training epochs, input sequence length, and score cutoff were determined by 5-fold cross validation (5CV).

Following tuning and cross-validation, we observed the optimal mean 5CV performance of 0.754 Spearman correlation and 0.751 Pearson correlation (Fig. S1). We noted that this new model showed comparable in the the held-out 20-percent test set, with a 0.746 Spearman correlation and 0.744 Pearson correlation (Fig. 3A).

**Figure 3.**
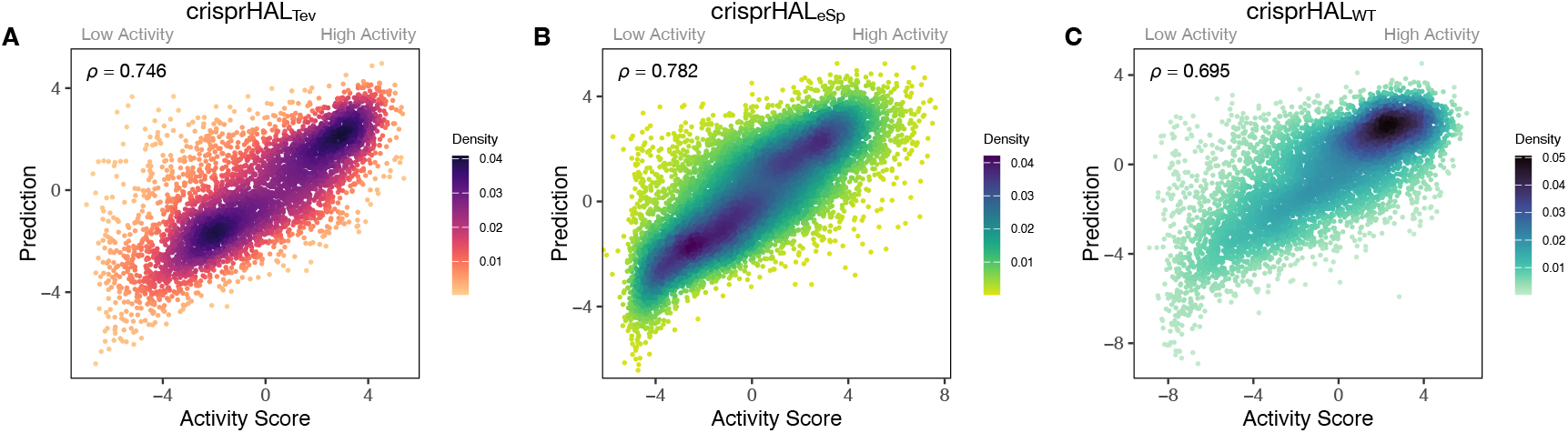
Predictions versus actual log_2_FC activity scores for each model’s respective 20% held out test set. (A)crisprHAL_Tev_ on the *C. rodentium* TevSpCas9 testing set of n=4253, (B) crisprHAL_eSp_ on the *E. coli* eSpCas9 testing set of n=11898, and (C) crisprHAL_WT_ on the *E. coli* SpCas9 testing set of n=6607.

### 3.2 Curation of prior eSpCas9 and SpCas9 data for better models

We now show how strategy used for the optimized crisprHAL_Tev_ model is generally useful when using datasets obtained from the literature. To provide maximal opportunity for our model to learn sequence features associated with eSpCas9 activity, we chose to regenerate the dataset without the sgRNA inclusion criteria present in the existing eSpCas9 dataset from Guo et al. (2018). From an initial pool of 64021 sgRNAs, using a minimum average control condition read count criteria of 31 (Table 2, and described in later sections), we obtain a final eSpCas9 dataset of 59489 sgRNAs and corresponding scores — 14419 more than the original version of the dataset from Guo et al. (2018). The sgRNAs not present in the original eSpCas9 dataset are distributed across a similar range of scores to the pre-existing sgRNAs, however, do cluster more towards lower activity (Fig. S2). This expanded dataset is then split into 80-percent training (n=47591) and 20-percent testing (n=11898) sets, with the training set used for all subsequent eSpCas9 model 5-fold cross validation testing (Table 3).

Our eSpCas9 model, crisprHAL_eSp_ shows improved performance versus the existing eSpCas9 models with a Spearman correlation of 0.783 and a Pearson correlation of 0.780 (Fig. 3B) in the held-out data. This performance is obtained following 62 epochs as determined through 5-fold cross validation. The best of the existing eSpCas9 models, DeepSgRNA_eSp_. reports a mean 5-fold cross validation Spearman correlation correlation of 0.715, which is substantially lower than the value the new model shows with held-out data. The other eSpCas9 model, from Guo et al. (2018), obtains a Spearman correlation of 0.682 on its held out 20% test set. To highlight the importance of the data, using the original *E. coli* eSpCas9 dataset, Guo et al. (2018) with the 43nt input used by DeepSgRNA (Wang and Zhang, 2019) — we obtain a Spearman correlation of 0.717 and a Pearson correlation of 0.716 on a held out 20% test set (n=9015).

We also reconstructed a new dataset from the *E. coli* SpCas9 data from Guo et al. (2018) to support the proposed data curation methods discussed. Using an average control condition read count cutoff of 73 (Table 2, and later sections), we obtain 33034 sgRNAs and scores from an initial pool of 61002 — a reduction of 11129 sgRNAs from original version of the dataset from Guo et al. (2018). Following this curation, the *E. coli* SpCas9 dataset was split into an 80-percent training (n=26427) and 20-percent testing (n=6607) sets (Table 3).

After training, the *E. coli* SpCas9 model, crisprHAL_WT_, shows a Spearman correlation and Pearson correlation of 0.695 and 0.720 on the held out test set (Fig. 3C). The Guo_WT_ and DeepSgRNA_WT_ models, constructed on the prior *E. coli* SpCas9 data, obtain 0.542 and 0.582 Spearman correlation — a difference of 0.153 and 0.113 respectively. To provide a direct comparison, akin to the eSpCas9 test, training the crisprHAL model using the original *E. coli* SpCas9 dataset results in a peak performance of 0.582 Spearman correlation and 0.611 Pearson correlation on their respective held out 20% test set (n=8833). To emphasize the importance of curation, the original model obtains substantially lower performance when using the full-size dataset despite training on 33% more data than does the new model trained on the curated dataset.

### 3.3 Limiting dynamic range inhibits performance

In this section we show the importance of choosing an appropriate minimum read-count cutoff for each dataset that maximizes the performance of the model trained on that curated dataset. Model training is reliant upon the accuracy of the data itself, both the input feature and label; i.e., the target site sequence and the activity score. The log_2_FC scores are reliant upon sufficient read count abundance in the control condition to facilitate a robust dynamic range in scores for model use. When we examine the entire set of *C. rodentium* TevSpCas9 sgRNA activity scores versus their control condition read counts we note a substantial score reduction associated with low control condition read counts (Fig. 4A). Since these data are generated using a depletion assay, low control condition read counts are expected to reduce the range of scores obtainable for highly active sgRNA sites; for example, a control read count of 1 allows only a read count of 0 to result in a positive score if an absolute value-based difference between calculation is used. This reduction can be seen across both the *E. coli* eSpCas9 and SpCas9 activity scores as well (Fig. 4B-C). Given the potential for these low quality scores to impact model performance, we generated subsets of the *C. rodentium* TevSpCas9 dataset by incrementally increasing the control condition read-count minimum (Fig. 1A). Comparing a control read-count of 1 to the previously used minimum of 20 (Guo et al., 2018), we see a substantial improvement; a 5CV mean Spearman correlation increase of 0.036 (Fig. 4E). As we increase this minimum count cutoff we see a steady increase in performance, peaking at a Spearman correlation of 0.754 with a read-count cutoff of 56, shown in purple in (Fig. 4E). This cutoff removes 16.2% of the data; a loss of 4894 sgRNAs (Fig. 4D). Given the implication that including these guides inhibits model training, we next examined the error between log2FC scores and the predictions from crisprHAL_Tev_ for all sgRNAs eliminated by the 56 cutoff, separated by control read count (Fig. S3). As expected, we note a high mean absolute error for between-model predictions and sgRNAs with low control condition read counts. When error is separated by positive and negative predictions, the loss of predictive utility is observed to occur in the positive predictions. This is unsurprising, as low control condition read counts reduce the precision of the readout for potentially active on-target sites. Above control read counts of 10-to-20 the error rate is substantially reduced and slowly decreases, in line with the model performance versus control cutoff results.

**Figure 4.**
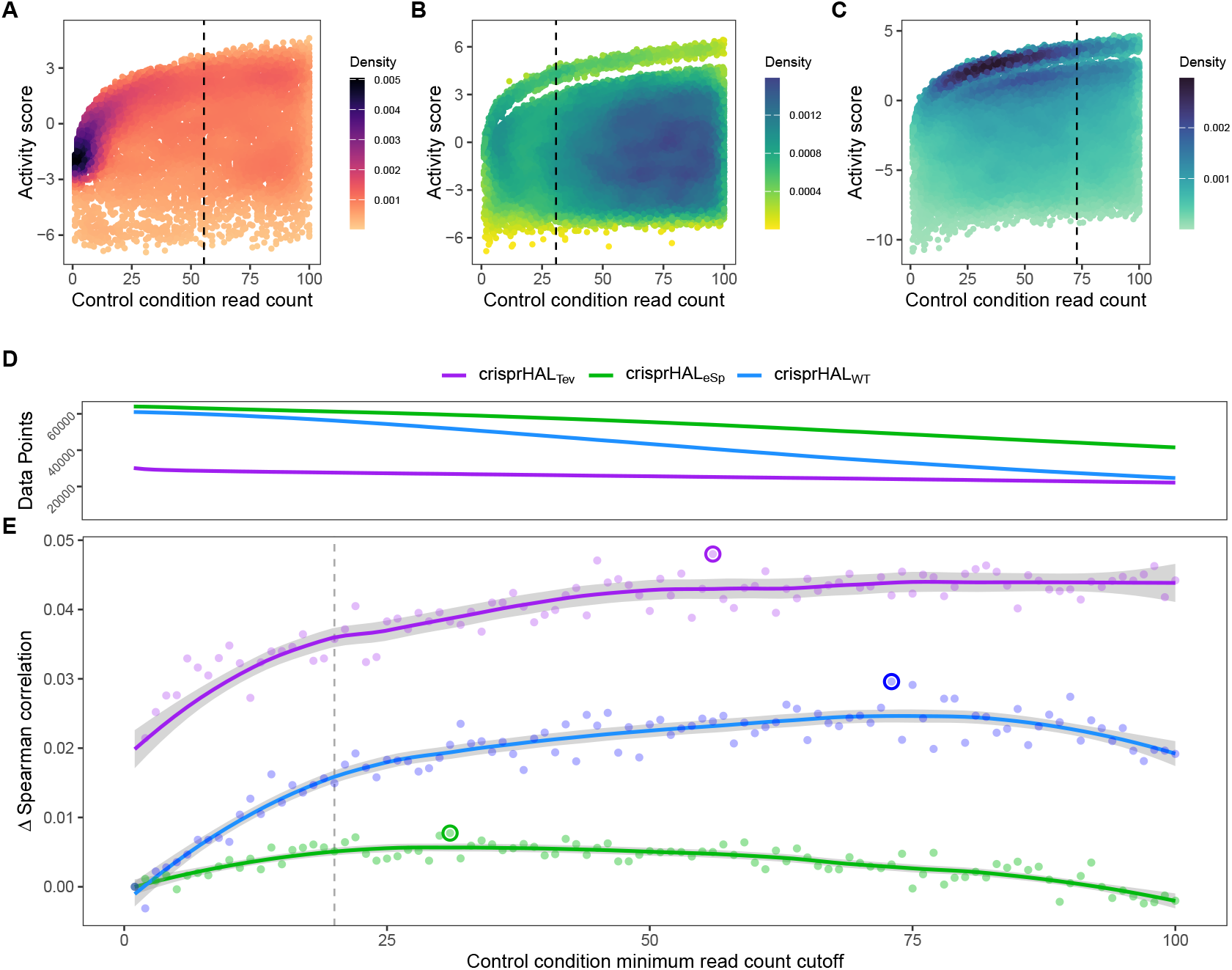
Control-condition read-counts and cutoff impacts on dataset size and model performance. The log_2_FC activity scores versus average control condition read counts from 1-to-100 for the (A) *C. rodentium* TevSpCas9, *E. coli* eSpCas9, and (C) *E. coli* SpCas9 datasets, each with their respective final model minimum read count cutoff. (D) Total data points in the *C. rodentium* TevSpCas9 (purple), *E. coli* eSpCas9 (green), and *E. coli* SpCas9 (blue) datasets at each control condition minimum read count. (E) Change in mean 5CV Spearman correlation values across increasing minimum control condition read counts for the *C. rodentium* TevSpCas9 model (purple), the *E. coli* eSpCas9 model (green), and the *E. coli* SpCas9 model (blue) versus a cutoff value of 1. The best performing cutoff value for each dataset and subsequent final model is circled. A previously used cutoff value of 20 used is indicated with the dashed line (Guo et al., 2018).

When we examine the impact of optimizing the control-condition read-count cutoff on the *E. coli* eSpCas9 dataset we again see an increase in model performance (green line in the figure), though not as substantial as seen as in the *C. rodentium* TevSpCas9 example (purple). Model performance improves consistently as low cutoff values increase, peaking at an average control condition read count cutoff of 31 reads across the two control replicates. The use of a read count minimum is likely less impactful to *E. coli* eSpCas9 model performance due to the higher read counts across the dataset, with comparatively few sgRNAs with low control read counts. Only 4.37% of sgRNAs in the eSpCas9 set have a mean control read count of less than 20 compared to 7.71% and 8.30% in the SpCas9 and TevSpCas9 sets, respectively. It would follow that datasets containing more sgRNAs with low quality activity scores will see a larger performance benefit from their removal.

When this process is applied to the *E. coli* SpCas9 dataset, we again observe a similar trend, with model performance improving until a minimum average control read count of 73 is reached. At this cutoff the 5CV average Spearman correlation is 0.697, an improvement of 0.029 over a cutoff of 1, and an improvement of 0.014 over using a cutoff of 20 (Fig. 4E). This performance boost is notable as 45.8% of the total data is removed in this example, reducing the dataset size from 61002 to 33034 sgRNA species. The loss of a large fraction of the dataset is unsurprising given that this dataset has the lowest read count coverage across all tested datasets. Thus we conclude that removing lower quality scores outweighs the benefit of retaining more training data across multiple datasets.

### 3.4 Adjacent nucleotides contain on-target activity information

We next examined the effect on predictive accuracy as a function of the model input sequence length. Prior work has noted the on-target Cas9 activity information contained in the nucleotides adjacent to the sgRNA target site, especially in the nucleotides immediately downstream of the PAM (Wang and Zhang, 2019; Ham et al., 2023). To test the performance benefit of including additional nucleotides upstream and downstream of the sgRNA target site, we train our TevSpCas9, eSpCas9, and SpCas9 models with input sequences of increasing length. We started from just the sgRNA and increased the sequence length incrementally one nucleotide at a time either upstream or downstream, but not both, with performance measured by average 5-fold cross validation Spearman correlation (Fig. 1B).

Across all three models, the most substantial boost to performance was obtained when the nucleotides immediately downstream of the PAM were included in the input sequence (Fig. 5A). For crisprHAL_Tev_ this region extends 11 nucleotides downstream of the PAM, before the performance benefit of adding more nucleotides to the input sequence begins to level off. This trend can be seen when examining average log_2_FC activity scores for each di-nucleotide position within and surrounding the sgRNA target, where most information is located between the sgRNA, PAM, and 11 nucleotides downstream (Fig. 5B-D). Analysis of more distant positional preferences for TevSpCas9 reveals little information, in line with model performance (Fig. S4). We note that the *E. coli* SpCas9 model obtains the greatest performance boost from this downstream nucleotide inclusion, with a 5CV average increase of +0.07 Spearman correlation units when including the PAM plus 10 nucleotides downstream compared to just using the sgRNA (Fig. 5A). Though the *C. rodentium* TevSpCas9 and *E. coli* eSpCas9 models do also see a performance benefit, more information appears to be contained within the sgRNA target site itself, especially in the PAM-distal region.

**Figure 5.**
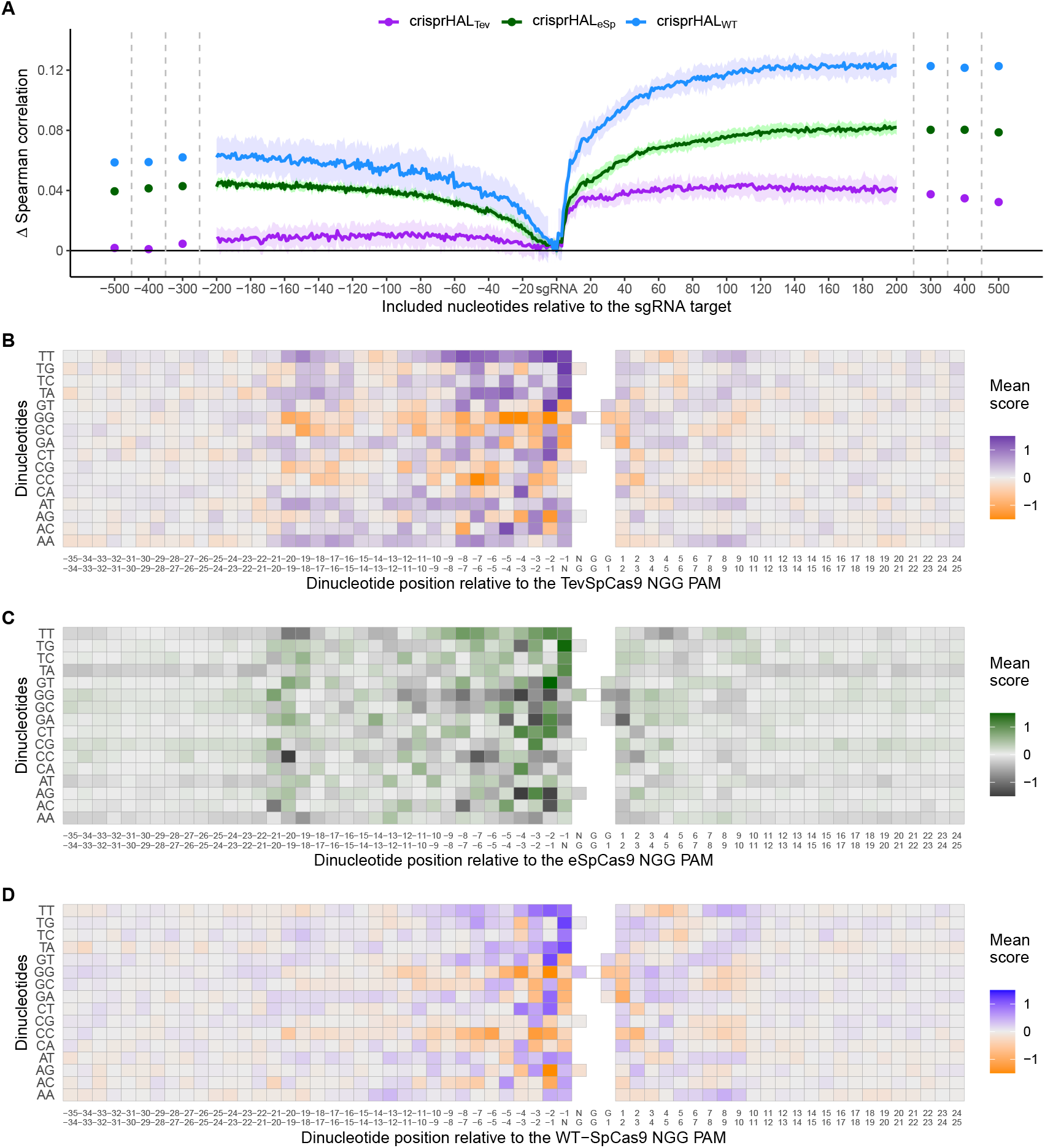
Target site and adjacent nucleotide on-target Cas9 activity information. (A) Model performance when upstream or downstream nucleotides are included alongside the sgRNA sequence as the model input. The change in model performance is reported as 5-fold cross validation mean Spearman correlation compared to a model input of just the sgRNA for *C. rodentium* TevSpCas9 and crisprHAL_Tev_ (purple), *E. coli* eSpCas9 and crisprHAL_eSp_ (green), and *E. coli* SpCas9 and crisprHAL_WT_ (blue). Mean log_2_FC score for each di-nucleotide option relative to the PAM for the (B) *C. rodentium* TevSpCas9, (C) *E. coli* eSpCas9, and (D) *E. coli* SpCas9 datasets.

Examining upstream nucleotides, there is a marginal performance benefit for crisprHAL_Tev_ when including 3 nucleotides, with a potential case to be made for including more. However, upon examination of the positional di-nucleotide mean scores across the *C. rodentium* TevSpCas9 and *E. coli* SpCas9 datasets, little activity information appears to be contained in this region (Fig. 5B/D). By contrast, we notice a trend within the *E. coli* eSpCas9 dataset which is present both upstream of the sgRNA target and beyond 10 nucleotides downstream of the PAM: A-T rich regions are disfavoured (Fig. 5C). Extending the di-nucleotide mean score plots 100 nucleotides upstream and downstream of the sgRNA-PAM site, this trend is maintained (Fig. S5). Though A-T rich adjacent regions are disfavoured, target sites with T rich PAM-proximal regions are very strongly favoured. This PAM-proximal T-preference is present across all three nucleases. There is also a G-preference for the N-position of the NGG PAM, in line with prior work suggesting local sliding of SpCas9 across adjacent PAM motifs (Corsi et al., 2022).

An interesting finding is the use of very long sequence length inputs for *E. coli* SpCas9 model performance. Appending 189 nucleotides upstream to the sgRNA target model input increases the mean 5CV Spearman correlation by 0.065, while appending the PAM plus 166 nucleotides downstream increases performance by 0.125 (Fig. 5A). Combined, a model built with a 378 nucleotide input sequence comprising 189 upstream, 20 sgRNA target, 3 PAM, and 166 downstream nucleotides obtains a mean 5CV Spearman correlation of 0.695, over 0.141 the performance seen when using an input of just the sgRNA-target. The sgRNA-target only model attains a mean 5CV Spearman correlation of 0.554. This finding is also repeated with the *E. coli* eSpCas9 model, with a moderate boost to performance seen when including 193 upstream and the PAM plus 190 downstream nucleotides — a total sequence length of 406. However, for eSpCas9, this can be more easily understood through the disfavouring of A-T rich regions, an attribute not seeming to be present for SpCas9 (Fig. S6).

### 3.5 SpCas9 model predictions generalize

Finally, we tested whether the models predicted on completely independent datasets collected using enrichment experiments rather than the depletion experiments from which the training sets were derived. We benchmarked crisprHAL_Tev_ against existing ML models using three independent enrichment datasets: *E. coli* pTox plasmid TevSpCas9 (n=260), pTox SpCas9 (n=255), and KatG TevSpCas9 (n=268) (Table 2). Across the three testing sets, using both Spearman and Pearson correlation metrics, crisprHAL_Tev_ achieves better performance than all 6 prior models tested — the original crisprHAL models, crisprHAL_Tev_ and crisprHAL_WT_, Guo_WT_, and DeepSgRNA_WT_ Cas9 models (Fig. 6). Specifically, the performance metrics for crisprHAL_Tev_ are: pTox TevSpCas9 0.690 Spearman correlation and 0.693 Pearson correlation, pTox SpCas9 0.650 Spearman correlation and 0.659 Pearson correlation, and KatG TevSpCas9 0.694 Spearman correlation and 0.615 Pearson correlation (Fig. 6A-C). Since the original crisprHAL models are constructed with the pTox TevSpCas9 and pTox SpCas9 datasets, only their performance on the KatG dataset is used for comparison. Performance from crisprHAL_eSp_ on the test sets is reported in Fig. S7 due to differences in enzyme preferences. Since SpCas9 and TevSpCas9 do not appear to share the characteristic of eSpCas9 that disfavours of AT-rich adjacent regions, the inclusion of such regions in crisprHAL_eSp_ reduces performance on the non-eSpCas9 datasets. When shorter input sequences from other models are used, that exclude the disfavoured AT-rich regions, crisprHAL_eSp_ performance on the TevSpCas9 and SpCas9 datasets improves (Fig. S7). We note that although the crisprHAL_Tev_ model is trained on a dataset which tests the TevSpCas9 nuclease, it still outperforms the models constructed on SpCas9 data on the pTox SpCas9 dataset, supporting the use of a TevSpCas9 data based model for SpCas9 prediction tasks.

**Figure 6.**
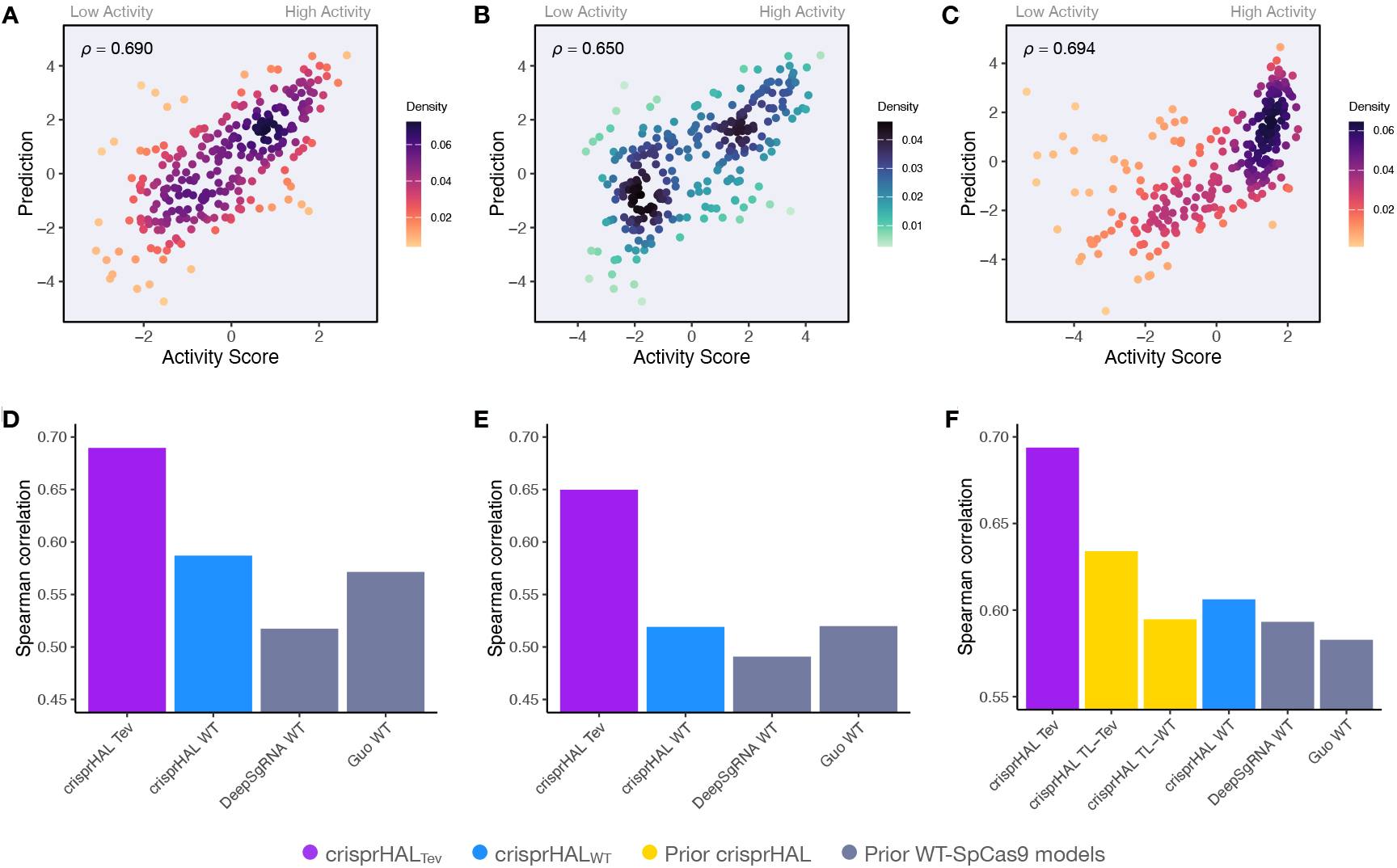
Benchmarking of the crisprHAL_Tev_ and crisprHAL_WT_ models. Predictions from crisprHAL_Tev_ trained on *C. rodentium* TevSpCas9 versus the log_2_FC activity scores for (A) the *E. coli* pTox plasmid TevSpCas9 activity set (n=260), (B) the *E. coli* pTox plasmid SpCas9 activity set (n=255), and (C) the *S. enterica* KatG target in *E. coli* TevSpCas9 activity set (n=268). Model performance comparisons of crisprHAL_Tev_ (purple) and crisprHAL_WT_ (blue), the original crisprHAL_TL-Tev_ and crisprHAL_TL-WT_ models (gold), and the prior models, DeepSgRNA_WT_ and Guo_WT_ (grey) on (D) the *E. coli* pTox plasmid TevSpCas9 activity set, (E) the *E. coli* pTox plasmid SpCas9 activity set, and **F)** the *S. enterica* KatG target in *E. coli* TevSpCas9 activity set.

A secondary result is the generalization of performance seen from crisprHAL_WT_ compared to both prior models constructed on the original *E. coli* SpCas9 data (Fig. 6D-F). Performance on the three testing sets exceeds both prior models, with the exception of pTox SpCas9, where performance is similar to the Guo_WT_ model. Using Pearson correlation as test set performance metric, similar results are attained (Fig. S8). We note that this performance is obtained using the ultra long 378 nucleotide model inputs.

## 4 DISCUSSION

In this paper we show the importance of data quality by examining the input features (target sites) and labels (activity scores) and develop best-in-class models for bacterial SpCas9 and eSpCas9 activity prediction. While a great deal of work has been spent on optimizing model architectures for CRISPR-Cas9 activity prediction, the model itself is only one part of the equation. Our work shows that a focus on curating the data from which the models are trained is as impactful on model performance as is developing novel ML approaches for this task.

The performance of a predictive model is constrained by its training data; and here we show that the accuracy of that data is as important as the size of the training set since smaller curated datasets can produce more accurate models than does the entire uncurated dataset. Inaccuracies with the feature (input) or label (score) can easily lead to sizable reductions in performance, and so data curation must be performed. Curation discards a small portion of a dataset due to low quality and ensures that the data are useful and accurate. However, as ML relies on large datasets, it is not obvious that setting a cutoff that eliminates nearly half of data points, as done with the *E. coli* SpCas9 data, would be expected to improve predictions. All three final models presented use minimum control read count cutoff values higher than previously used, with the SpCas9 and TevSpCas9 datasets seeing optimal performance in spite of the removal of substantial amounts of training data. Thus, increasing data quality outweighs reducing dataset size even when that reduction in data size is substantial. We found that the need for a cutoff arises from sparse control-condition read-counts that are found in depletion experiments. Very low read-counts do not provide an accurate estimate of the initial feature abundance, and so reduce the dynamic range of activity scores and their accuracy (Rapaport et al., 2013; Fernandes et al., 2013; Love et al., 2014). Alongside methods like the control read count cutoff discussed, other strategies to mitigate this issue include increasing sequencing depth to directly improve read count abundance, and using more control replicates to better represent the measurement variance.

Akin to score curation, little emphasis has been placed on optimizing model inputs for the inclusion of relevant target-site adjacent nucleotides. Some approaches across the landscape have been quite conservative, with the inclusion of only the sgRNA and sometimes the NGG PAM, missing out on substantial relevant information. The sgRNA target site, and more specifically the PAM-proximal side, does contain the most relevant information for on-target activity. This is in line with optimal model performance found by Ham et al. (2023) during their input sequence length optimization, as well as prior biological findings of downstream Cas9-DNA interactions Zhang et al. (2019); Yang et al. (2021). Across all three depletion datasets for TevSpCas9, eSpCas9, and SpCas9, we note substantial on-target information contained in the region encompassing the PAM and up to 11 nucleotides downstream. Of note is that nearly identical positional and di-nucleotide preferences can be seen across the three nucleases. Moving upstream from the sgRNA, limited arguments can be made for the importance of the 1-to-3 adjacent nucleotides. However, what is clear from the *E. coli* eSpCas9 data is a distinct disfavouring of AT rich target site adjacent regions, a preference extending beyond 100 nucleotides both upstream and downstream of the sgRNA — excluding the PAM and directly adjacent downstream nucleotides. A less obvious preference exists in the *E. coli* SpCas9 dataset, however no such notable trend occurs for the *C. rodentium* TevSpCas9 dataset. Given the overlap in sgRNA target sites between both *E. coli* datasets, it is unclear if the trend is due to enzymatic preferences, is species/strain specific, or if it is an artifact of sgRNA target site selection itself. However, what is obvious is the improvement to both held-out and independent test set performance when the *E. coli* SpCas9 model uses long input sequences containing nearly 200 nucleotides upstream and downstream of the target site. The long input sequences provide relevant information. Were this improvement only present within a single dataset, it would be easy to dismiss the improvement as being related to some aspect of assay design or another dataset characteristic. However, since the trend generalizes across datasets, distant nucleotides from the target site may impact SpCas9 activity.

When optimizations to input sequence length and score are applied, we obtain excellent performance on the three independent datasets from both the crisprHAL_Tev_ and crisprHAL_WT_ models. Though crisprHAL_Tev_ performs best for SpCas9 and TevSpCas9 activity prediction, and is the model recommended for such tasks, crisprHAL_WT_ is arguably more informative. Unlike the *E. coli* eSpCas9 dataset, when curation is performed for the SpCas9 dataset the amount of data remaining is reduced to a point lower than the original *E. coli* SpCas9 dataset. It is this reduction that makes performance on the held-out test set and, more importantly, the three independent test sets, so important. The *E. coli* SpCas9 model performs better, and more consistently, than the prior models on each test in spite of having less training data. Though crisprHAL_eSp_ does also boast improved performance compared to models trained on the original *E. coli* eSpCas9 data, the lack of independent testing data and the increased amount of training data limits strong statements about data curation-based performance improvement. However, what is notable is that when training crisprHAL with either original *E. coli* dataset, we obtain performance barely exceeding that of DeepSgRNA_eSp_. Though the crisprHAL architecture performs well for transfer learning applications, as it was designed to do, in this case it is not the architecture that is responsible for the performance improvements; it is the data.

Here we present crisprHAL_Tev_, a deep learning model designed for bacterial sgRNA/SpCas9 prediction tasks with performance that exceeds that of existing models across all tested datasets. This performance was obtained through focusing on the quality of the model inputs and scores rather than through the common method of a novel architecture or other machine learning methodology. To support this data-centred approach, we revisited two prior datasets measuring eSpCas9 and SpCas9 activity in *E. coli*, remaking the data, and generating subsequent models with improved performance. During analysis of the model inputs we noted a trend of identical downstream nucleotide on-target activity information which suggests that the nucleotides directly downstream of the PAM have an notable impact on activity, with more distant upstream and downstream nucleotides also seeming to impact activity. Using these methods we present models which provide best-in-class SpCas9 and eSpCas9 predictions and demonstrate the importance of data curation to improve model performance.

## 5 CONCLUSION

A great deal of work has been spent on the optimization of model architectures for CRISPR-Cas9 activity prediction, however the model itself is only one part of the equation. Across the sgRNA/Cas9 prediction field, numerous papers have been published which present novel machine learning approaches and updates to improve activity prediction. While these papers succeed at providing performance improvements, relatively little focus has been spent on cleaning and optimizing the data from which the models are trained. Here we show the importance of examining the target site inputs and activity score labels through the development of best-in-class models for bacterial SpCas9 and eSpCas9 activity prediction.

## 6 ADDITIONAL INFORMATION AND DECLARATIONS

### 6.1 Competing Interests

The authors declare that they have no competing interests.

### 6.2 Author Contributions

Tyler S. Browne conceived and designed the experiments, performed the computational work, analyzed the data, prepared figures and/or tables, authored or reviewed drafts of the article, and approved the final draft. David R. Edgell conceived and designed the experiments, authored or reviewed drafts of the article, and approved the final draft. Gregory B. Gloor conceived and designed the experiments, authored or reviewed drafts of the article, and approved the final draft.

### 6.3 Data Availability

The prediction model series crisprHAL is available at: https://github.com/tbrowne/crisprHAL. To reproduce the presented figures and exact model performance, please use the following repository: https://github.com/tbrowne5/Better-data-for-better-predictions-data-curation-improves-deep-learning-for-sgRNA-Cas9-prediction. The latter repository also contains raw data and other relevant code for data processing and dataset generation.

### 6.4 Funding

This work was supported by the Canadian Institutes of Health Research (PJT-191939 and PJT-159708) National Sciences and Engineering Research Council of Canada (RGPIN-06519-2025), the Ontario Graduate Scholarship (Tyler S. Browne), and the Queen Elizabeth II Graduate Scholarship in Science and Technology (Tyler S. Browne).

## ACKNOWLEDGEMENTS

We would like to thank Michael T. Hallett and Dalton T. Ham for their support through discussions on machine learning, and sgRNA/Cas9 molecular biology, respectively.

## Notes

https://github.com/tbrowne5/Better-data-for-better-predictions-data-curation-improves-deep-learning-for-sgRNA-Cas9-prediction

https://github.com/tbrowne5/crisprHAL

### Competing Interest Statement

The authors have declared no competing interest.

