## Supplementary material for "Better data for better predictions: data curation improves deep learning for sgRNA/Cas9 prediction": Fig. S

### SUPPLEMENTARY FIGURES


**S1** Cross validation performance for crisprHAL$_{Tev}$, crisprHAL$_{eSp}$, and crisprHAL$_{WT}$ across tested training epochs

**S2** eSpCas9 and SpCas9 log$_2$FC score distributions for original and curated dataset versions

**S3** crisprHAL$_{Tev}$ model predictions of *C. rodentium* TevSpCas9 data removed by the minimum control read count cutoff of 56

**S4** Upstream and downstream di-nucleotide mean score plots for the *C. rodentium* TevSpCas9 dataset

**S5** Upstream and downstream di-nucleotide mean score plots for the *E. coli* eSpCas9 dataset

**S6** Upstream and downstream di-nucleotide mean score plots for the *E. coli* wild-type SpCas9 dataset

**S7** crisprHAL$_{eSp}$ model performance on the hold-out and independent test sets using different input sequence lengths

**S8** Pearson correlation metrics for crisprHAL$_{Tev}$, crisprHAL$_{WT}$, and prior models on the 3 independent test sets

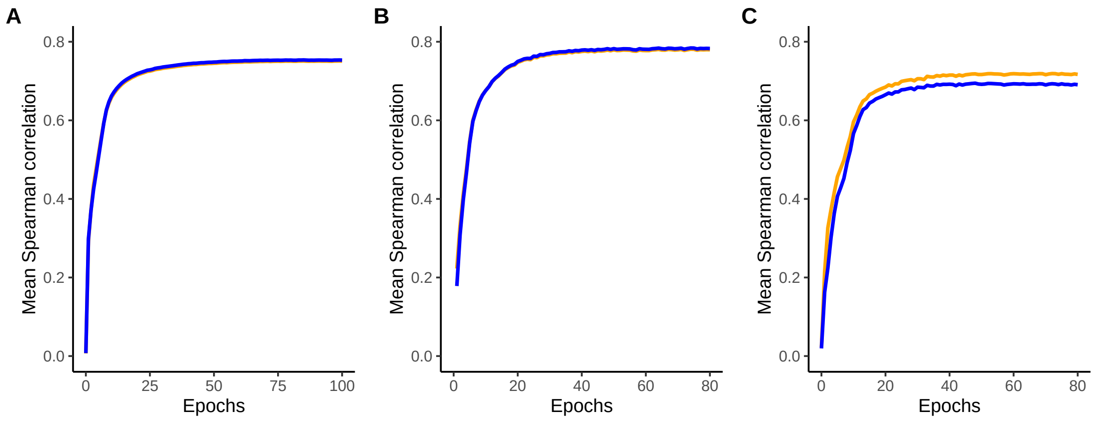


**Figure S1.** Mean 5-fold cross validation Spearman correlations (blue) and Pearson correlations (orange) for (A) crisprHAL$_{\mathrm{Tev}}$, (B) crisprHAL$_{\mathrm{eSp}}$, and (C) crisprHAL$_{\mathrm{WT}}$ across tested training epochs. The epoch providing the best mean Spearman correlation for each respective model is used for final model training.

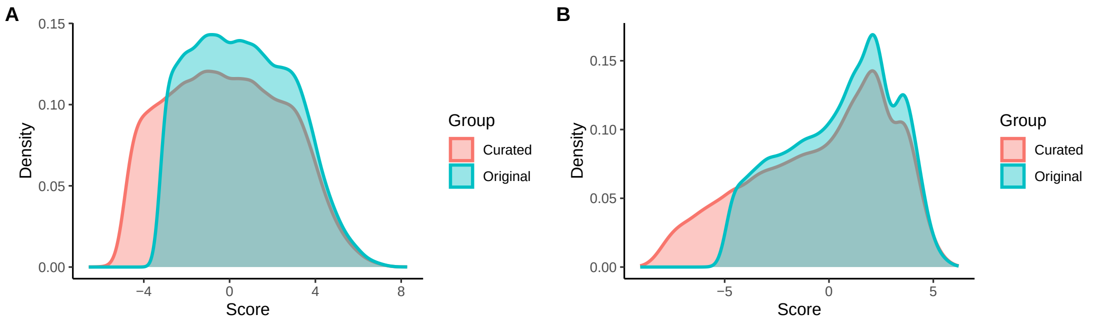


**Figure S2.** Densities of $\log_2$FC scores for the (A) *E. coli* eSpCas9 dataset and (B) *E. coli* wild-type SpCas9 dataset. The densities in orange show the distribution of the scores for each dataset following our data curation, with densities in blue showing the distribution of scores for data points in the original, prior, version of each dataset.

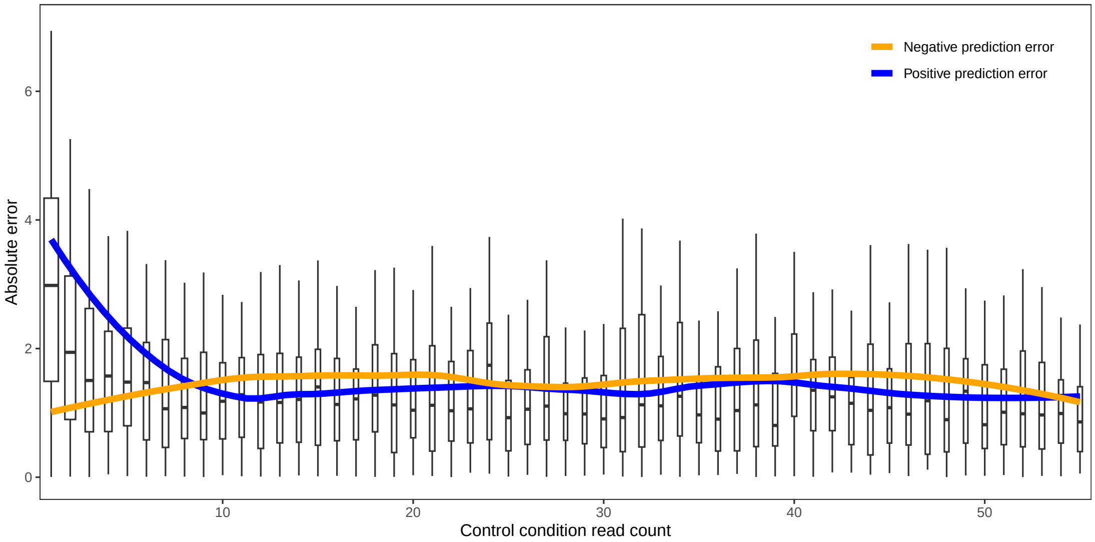


**Figure S3.** Error between predictions from crisprHAL$_{Tev}$ and the *C. rodentium* TevSpCas9 data removed by the optimal identified minimum control condition read count cutoff of 56. Error between predictions and $\log_2$FC scores is separated by control condition read count. Trend lines show mean absolute error between predictions and positive (blue) and negative (orange) $\log_2$FC scores.

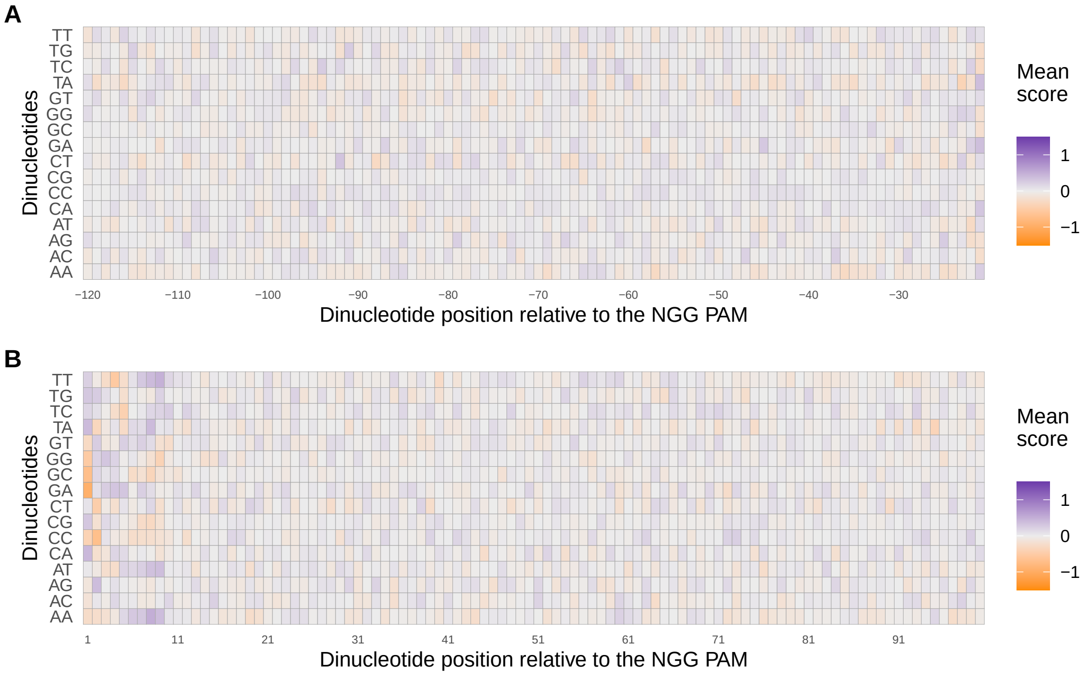


**Figure S4.** (A) Upstream and (B) downstream mean $\log_2$FC scores from the *C. rodentium* TevSpCas9 dataset for each di-nucleotide option adjacent to the target site and NGG PAM. Positions are labelled with respect to moving upstream or downstream of the PAM.

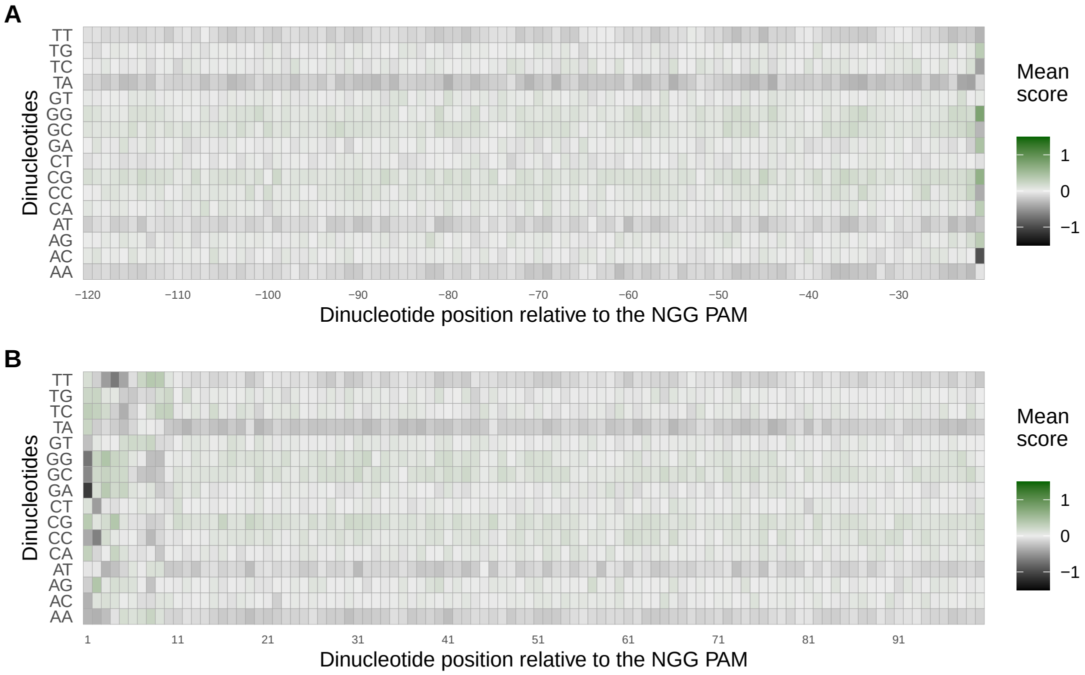


**Figure S5.** (A) Upstream and (B) downstream mean $\log_2$FC scores from the *E. coli* eSpCas9 dataset for each di-nucleotide option adjacent to the target site and NGG PAM. Positions are labelled with respect to moving upstream or downstream of the PAM. Of note is the consistent disfavouring of AT-rich target site adjacent regions by eSpCas9.

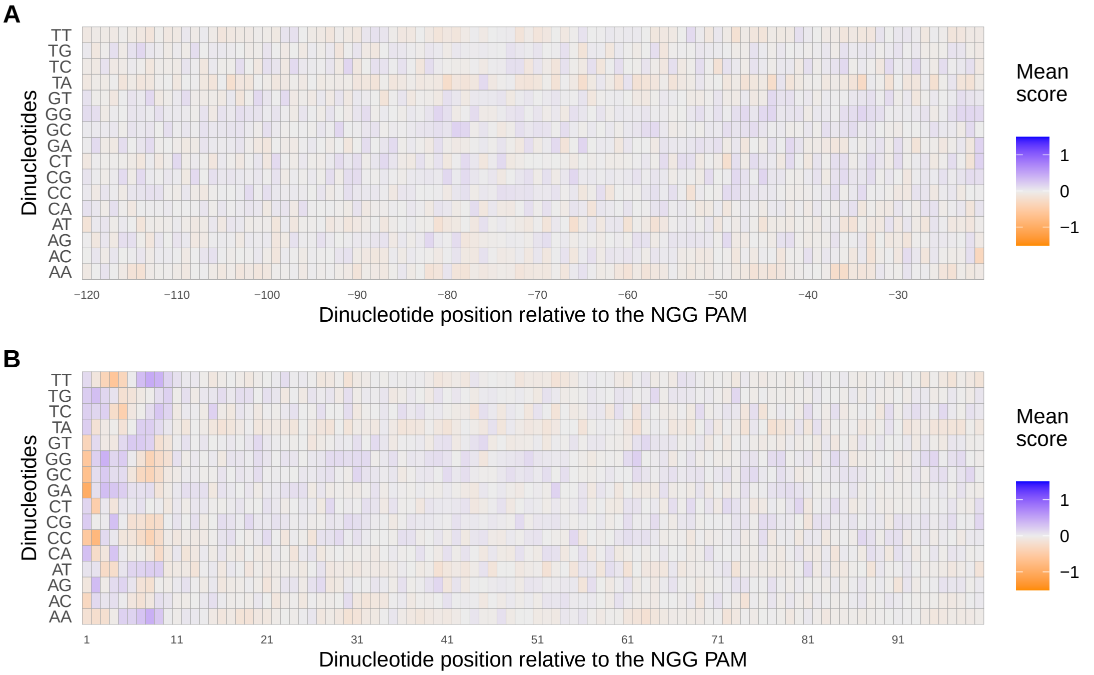


**Figure S6.** (A) Upstream and (B) downstream mean $log_2FC$ scores from the *E. coli* wild-type SpCas9 dataset for each di-nucleotide option adjacent to the target site and NGG PAM. Positions are labelled with respect to moving upstream or downstream of the PAM.

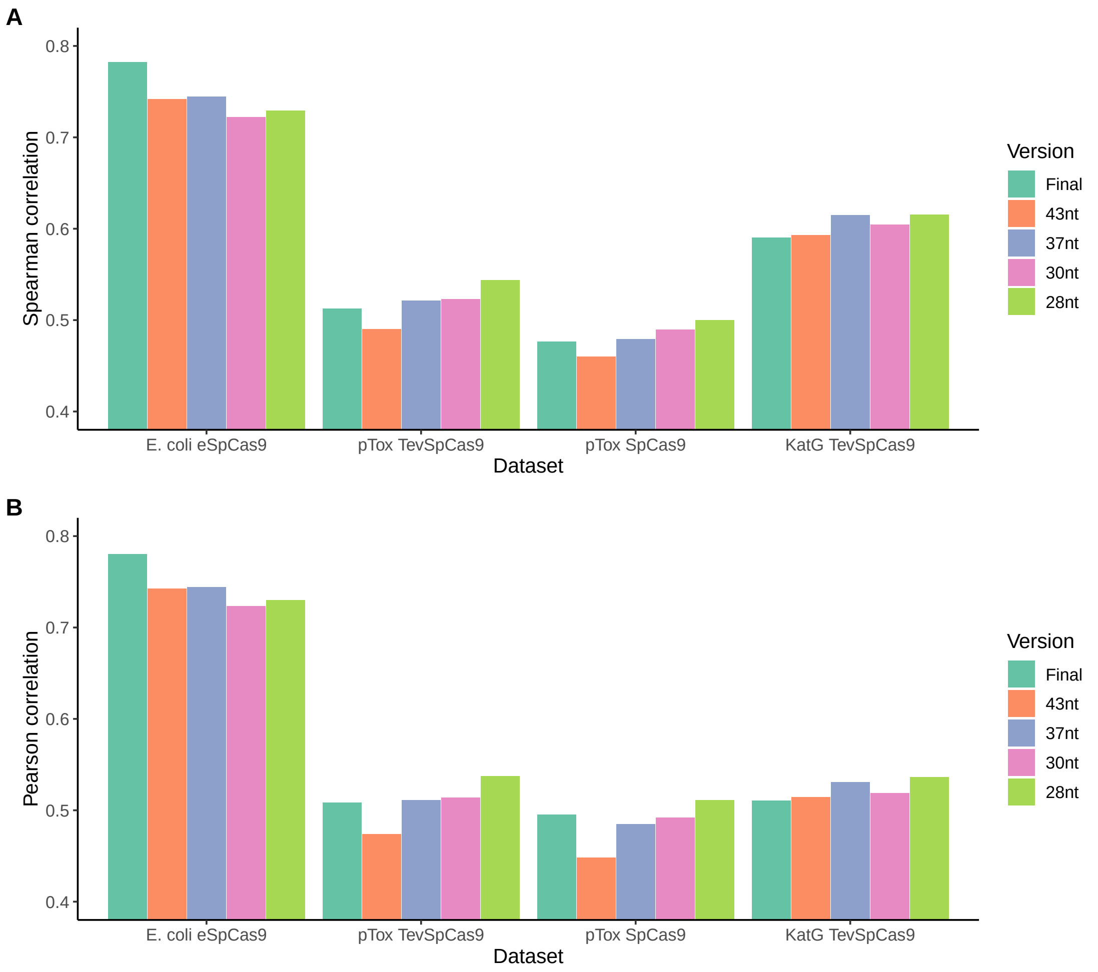


**Figure S7.** Performance from crisprHAL$_{eSp}$ model on the hold-out and independent test sets when using 5 different input sequence lengths, as measured by (A) Spearman correlation, and (B) Pearson correlation. The inputs tested with adjacent nucleotide inclusion (U=upstream, D=downstream) are: the final crisprHAL$_{eSp}$ 406 nt input (U=193, D=193), 43 nt used for DeepSgRNA$_{eSp}$ (U=10, D=10), 37 nt used for crisprHAL$_{Tev}$ (U=3, D=14), 30nt used for Guo$_{eSp}$ (U=4, D=6), and 28 nt used for crisprHAL$_{TL\text{-}Tev}$ and crisprHAL$_{TL\text{-}WT}$ (U=0, D=8). Given the unique target site adjacent nucleotide preferences by eSpCas9, the inclusion of these regions in the model input improves eSpCas9 performance, but hinders performance on wild-type SpCas9 and TevSpCas9 tasks. When the upstream nucleotides, and most downstream nucleotides, are excluded from the input, as per the 28 nt input sequence length, performance on the wild-type SpCas9 and TevSpCas9 test sets is improved.

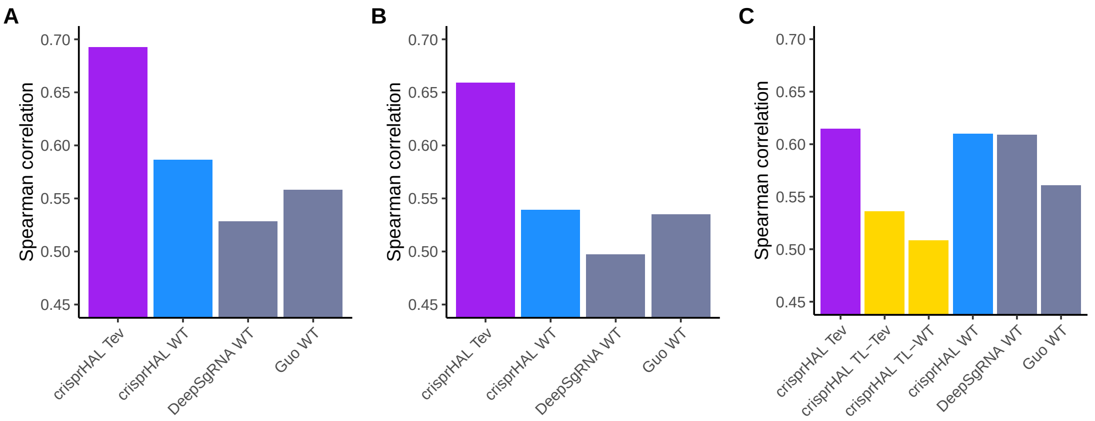


**Figure S8.** Pearson correlation model performance comparisons of crisprHAL$_{\text{Tev}}$ (purple) and crisprHAL$_{\text{WT}}$ (blue), the original crisprHAL$_{\text{TL-Tev}}$ and crisprHAL$_{\text{TL-WT}}$ models (gold), and the prior models, DeepSgRNA$_{\text{WT}}$ and Guo$_{\text{WT}}$ (grey) on (A) the *E. coli* pTox plasmid TevSpCas9 activity set, (B) the *E. coli* pTox plasmid SpCas9 activity set, and (C) the *S. enterica* KatG target in *E. coli* TevSpCas9 activity set.
